# Development of 2-phenylethanol plus acetic acid lures to monitor obliquebanded leafroller (Lepidoptera: Tortricidae) under mating disruption

**DOI:** 10.1101/093955

**Authors:** A. L. Knight, G. J. R. Judd, E. Basoalto, A. M. El-Sayed

## Abstract

We evaluated the effectiveness of 2-phenylethanol (PET) in combination with acetic acid (AA) as a binary lure for monitoring male and female obliquebanded leafroller, *Choristoneura rosaceana* (Harris). Studies were conducted in apple, *Malus domestica* Borkhausen, orchards treated with or without sex pheromone dispensers for mating disruption (MD). Open polypropylene vials, closed membrane cups, and rubber septa loaded with AA and/or PET in varying amounts were first evaluated in a series of trapping experiments. Membrane cups loaded with 800 mg of PET were as effective as 10-mg septa, but longer lasting, and were comparable to the open vials. A membrane cup AA lure was effective in tests, but further work is needed to increase its release rate and extend its activity. Catches of codling moth, *Cydia pomonella* (L.), and *C. rosaceana* were unaffected by combining PET with (*E*,*E*)-8,10-dodecadien-1-ol, the sex pheromone of codling moth, pear ester, (*E*,*Z*)-2,4-ethyl-decadienoate, and AA lures. Adding (*E*)-4,8-dimethyl-1,3,7-nonatriene to this blend to enhance codling moth catch, significantly reduced catches of *C. rosaceana.* PET+AA was a more attractive binary lure than AA plus phenylacetonitrile (PAN) for *C. rosaceana.* The addition of PET or PAN to traps already baited with the sex pheromone of *C. rosaceana* significantly reduced male catches. Traps baited with PET+AA placed in blocks not treated with MD caught significantly fewer *C. rosaceana* than traps baited with sex pheromone. In comparison, sex pheromone-baited traps in MD blocks caught ≤ 1 male moth per season which was significantly lower than total moth (> 10) or female moth (≥ 3) catch in these blocks with PET+AA. A high proportion (> 70%) of trapped females were mated in both untreated and MD-treated orchards. Further refinement of this binary, bisexual lure using membrane cup technology may allow the establishment of action thresholds and improve management timings for *C. rosaceana*.

## Introduction

Commercial apple, *Malus domestica* Borkhausen, orchards in western North America can be attacked by a suite of tortricid pests (leafrollers) that feed primarily on foliage but sometimes damage the surface of fruits (Madsen and Madsen 1980; Barnett et al. 1991; Beers et al. 1993). *Pandemis pyrusana* Kearfott, and the obliquebanded leafroller, *Choristoneura rosaceana* (Harris), are the two most important pests in this leafroller complex in Washington State, (Brunner and Beers 1990). *C. rosaceana* is also an important pest of apple in eastern North America (Chapman 1973). Improved monitoring of these pests is needed because catches of male tortricid leafrollers with sex pheromone-baited traps have not been effective in characterizing within-field population densities in apple orchards (Knight 2001; Alway 2003). The broad host range of these tortricid pests and their dispersal from outside habitats (via larval ballooning and adult flights) into management units are thought to contribute to the difficulty in using traps to monitor and predict local population densities (Chapman 1973; Thayer et al. 1997; Knight 2001).

Apple growers usually manage leafrollers with insecticides which has selected for resistance and cross-resistance to some classes of compounds (Pree et al. 2001; Dunley et al. 2006). In order to counter this selection pressure many growers have broadened their management tactics to include use of microbial and newer conventional insecticides (Cossentine et al. 2003; Smirle et al. 2003; Sial et al. 2010). Meanwhile, researchers have examined the use of sex pheromones for mating disruption (MD), including formulations targeting several species (Evenden et al. 1999; Curkovic and Brunner 2005; Judd and Gardiner 2008). At present, MD is being used against leafrollers on a relatively small proportion of apples in Washington State; primarily against *C. rosaceana* in certified-organic orchards (Knight 2010). Catches of *C. rosaceana* with sex pheromone traps are significantly reduced (>92%) in orchards treated with MD (Knight et al. 1998; Judd and Gardiner 2008). Thus, while sex pheromone-baited traps can be used to assess whether sex pheromone communication of adult populations is being affected, their catches are generally too low to track the seasonal phenology of these pests. Consequently, growers must rely on model predictions to time sprays (Jones et al. 2005) with an absence of knowledge of the relative pest population densities.

Recently, pear ester, ethyl (*E*, *Z*)-2,4-decadienoate plus acetic acid in traps with the sex pheromone of codling moth, *Cydia pomonella* (*L*.), (*E*,*E*)-8,10-dodecadien-1-ol (codlemone), was shown to be a mildly attractive lure for apple leafroller species common in Washington State (Knight et al. 2014). Surprisingly, the relatively low moth counts were well correlated with immature population densities of leafrollers within orchards, and thus were useful predictors to aid pest management decisions (Knight et al. 2014). Further development of more powerful non-pheromone attractants for leafrollers could improve the accuracy of this approach, especially in orchards under management with MD (Knight et. al 1998).

A number of common host-plant volatiles, including green leaf volatiles and terpenes, in combination with acetic acid, have been tested as potential lures for adult leafrollers in apple (Knight et al. 2014). In addition to pear ester, these have included (*E*)-β-farnesene, (*E*)-β-ocimene, (*E*)-4,8-dimethyl-1,3,7-nonatriene, (*Z*)-3-hexenyl acetate, and (*E*,*E*)-farnesol. Unfortunately, none of these volatiles significantly increased the catches of adult leafrollers achieved with using acetic acid alone.

The search for new attractants for leaf-feeding tortricids is being directed in part by the characterization of herbivore-induced plant volatiles (HIPVs) produced by apple leaves in response to feeding by two species of leafrollers: the light brown apple moth, *Epiphyas postvittana* (Walker) (Suckling et al. 2012), and the dark fruit tree tortrix, *Pandemis heparana* Denis and Schiffermüller (Giacomuzzi et al. 2013). Phenylacetonitrile was recently identified among a subset of these identified HIPVs as being an effective attractant for *P. pyrusana* in Washington State, but only when combined with acetic acid (Knight et al. 2016). Surprisingly, this binary lure caught six-times more total moths (1:1 sex ratio) than the number of male moths caught with a commercial sex pheromone lure. Another aromatic benzenoid volatile, 2- phenylethanol was not originally reported to be released from apple foliage injured by tortricid larvae, but has more recently been found to be released at low levels, and also found to be attractive for both sexes of a number of tortricid adults when combined with an acetic acid colure in Europe, New Zealand, and Canada (El-Sayed et al. 2016; Giacomuzzi et al. 2016).

The alcohol, 2-phenylethanol is a common compound usually synthesized by plants from phenylalanine (Bruce et al. 2005); and is found in a wide array of flowers and fruits, including apple (Omata et al. 1990; Knudsen et al. 1993; Buchbauer et al. 1993). In tree fruits, 2-phenylethanol is also a bisexual attractant for apple fruit moth, *Argyresthia conjugella* Zeller (Bengtsson et al. 2006), and an attractant for the green lacewing, *Chrysoperla carnea* (Say) (Zhu et al. 2005). Recognition of 2-phenylethanol as a HIPV is consistent with an earlier study by Boevé et al. (1996) with apples infested with the European apple sawfly, *Hoplocampa testudinea* (Klug). Also, 2-phenylethanol has previously been combined with acetic acid to enhance its attraction for *Drosophila melanogaster* Meigen, (considered a microbial volatile in this context) (Zhu et al. 2003; Becher et al. 2010). Acetic acid and 2-phenylethanol are part of a blend of volatiles released by grape inoculated with either yeasts or acetic bacteria that were shown to increase oviposition and fitness of the tortricid *Lobesia botrana* (Denis and Schiffmüller), (Tasin et al. 2011). Combinations of acetic acid and 2-phenylethanol have also been shown to increase catches of adult *Chrysoperla* spp., in apple and other tree fruit crops (Toth et al. 2009; Jones et al. 2015).

Following this line of research, we conducted studies to evaluate the effectiveness of using 2-phenylethanol in combination with acetic acid as a potential bisexual lure for adult *C. rosaceana* in orchards treated with and without MD. Parallel studies were conducted to develop effective long-lasting seasonal lures. In addition, three additional studies were conducted to evaluate the potential effectiveness of this binary lure in apple pest management: 1. the effectiveness of 2-phenylethanol versus phenylacetonitrile when both are used with acetic acid; 2. the use of either aromatic in combination with the sex pheromone of *C. rosaceana,* and 3. the use of 2-phenylethanol within the same trap with lures for *C. pomonella*.

## Materials and Methods

### Lure development

Several types of release devices were prepared with 2-phenylethanol (99%) and glacial acetic acid (99.7%) (Sigma Aldrich, St. Louis, MO). Four proprietary plastic membrane cup lures (1.8 cm diameter) loaded with acetic acid (TRE3321 and TRE1468), 2-phenylethanol (TRE1256), or phenylacetonitrile (TRE1381) were prepared by Trécé Inc. (Adair, OK). A 5-cm diameter membrane cup lure loaded with 2-phenylethanol (TRE1274) was also provided by Trécé Inc. All membrane cup lures were loaded with neat material.

A second release device consisted of polypropylene vials (Nalg-Nunc International, Rochester, NY) loaded with two cotton balls and with different sized apertures drilled into the cap. The 2-phenylethanol vial lures consisted of a 4-ml vial with a 1.5-mm aperture and loaded with 3-ml, and the two acetic acid vial lures tested were 8.0-ml vials with either 1.0 or 3.2-mm apertures and loaded with 5.0 ml. Red rubber septa lures with 2-phenylethanol were prepared using dichloromethane as the carrier and loaded with 1.0, 5.0, and 10.0 mg. Four proprietary septa lures (Trécé Inc.) were also included in our studies: red rubber septa loaded with the full proprietary rate sex pheromone of *C. rosaceana* (Pherocon OBLR/W, TRE3223) and a 10% loading (TRE1229), and grey halobutyl septa (Pherocon CM DA Combo, TRE3461) loaded with codlemone and pear ester and a second lure including the addition of 3.0 mg of (*E*)-4,8-dimethyl-1,3,7-nonatriene (DMNT, TRE1225).

The weight loss of all acetic acid and 2-phenylethanol lures (N = 5) was measured in the laboratory at 25°C. Lures were spaced 15 cm apart on paper-lined trays, aired for 24 h and weighed, and then re-weighed weekly for up to 12 weeks. It was not possible to estimate the weight loss of 4.0-ml vials with 1.5-mm apertures loaded with 2-phenylethanol due to a gradual weight gain by the lure over time.

### Study protocols

Orange delta traps with sticky inserts (Pherocon VI, Trécé Inc.) were used in most studies, but white plastic bucket traps with green lids (Multipher 1, Great Lakes IPM, Vestaburg, MI) were used in two experiments. Multipher traps were filled with 250 ml of propylene glycol (Integra Chemical Co., Kent, WA) to retain moths. Five or 10 replicates of each lure treatment were randomized and spaced 50 m apart and 30 m from the physical border of each orchard in each experiment. Traps were hung in the tree canopy at a 2.5 – 3.0 m height. All lures were placed in the center of the trap’s sticky liner (AlphaScents, Portland, OR).

Study sites (2 – 150 ha) included seven commercial and one unsprayed experimental orchard (Moxee, 46°30’5N, 120°10’16W). One orchard was certified organic (Azwell, 47°56’17’N, 122°44’7’W) and one was managed with a non-insecticide program (Ashland, 42°14’47’N, 119°52’22’W). The remaining orchards were managed with unspecified conventional spray programs. All of the orchards except Ashland were treated with sex pheromone dispensers for MD of *C. pomonella.* The Burbank orchard (46°19’51’N, 18°43’18’W) in 2015 and 2016 and the Azwell orchard in 2015 were treated with Isomate CM Flex (Pacific Biocontrol, Vancouver, WA) loaded with 88 mg of codlemone. The Azwell orchard in 2016 was treated with a dual aerosol system (SemiosBio Technologies, Vancouver, British Columbia) applied at 2.5 ha^−1^ that released 40 mg puffs (7.22 mg a.i.) of codlemone and (Z)-11-tetradecenyl acetate for *C. rosaceana* from separate canisters every 15 minutes for 12 hours each day. The Parker (46°29’25’N, 120°25’41’W), Wapato (46°24’33’N, 120°28’51’W), and Naches (46°39’56’N, 120°39’49’W) orchards in 2015 and the Moxee orchard in both 2015 and 2016 were treated with Cidetrak CMDA Combo PP dispensers loaded with 90 and 60 mg of codlemone and pear ester, respectively. The Wapato, Tieton (46°42’11’N, 120°45’21’W), and East Wenatchee (47°25’17’N, 120°17’17’W) orchards during 2016 were treated with Cidetrak CM-DA Meso dispensers loaded with 900 and 600 mg of codlemone and pear ester, respectively, and applied at 80 ha^−1^.

### 2015 lure studies

Three field studies were conducted in 2015 to compare moth catches of *C. rosaceana* with various combinations of acetic acid and 2-phenylethanol lures. Experiment 1 compared traps baited with 2-phenylethanol (4.0-ml vials with a 1.5-mm aperture) in combination with one of the three different acetic acid co-lures: the membrane cup TRE3321, and two polyethylene 8.0-ml vials with either 1.0 or 3.2-mm apertures. This study was conducted in the Burbank orchard from 14 - 28 July with 10 replicates per lure. Experiment 2 compared moth catches in traps baited with the 8.0-ml vial with the 3.2-mm aperture loaded with acetic acid and either the 4.0-ml vial with a 1.5-mm aperture or the TRE1256 membrane cup lure loaded with 2-phenylethanol. This test included five replicates and was conducted in the Burbank and Ashland orchards from 18 June to 2 July and from 31 July to 13 August, respectively. Experiment 3 compared moth catch in traps baited with acetic acid (8.0-ml vial with a 3.2-mm aperture) and either the smaller (TRE1256) or larger (TRE1274) membrane cup lures loaded with 2-phenylethanol. This was conducted with five replicates in the Burbank and Azwell orchards from 18 June to 2 July and 11 August to 8 September, respectively.

### 2016 lure studies

Four additional lure combination studies were conducted during 2016. Red septa lures (TRE1410, 1411, 1412) loaded with 1.0, 5.0, and 10.0 mg 2-phenylethanol, respectively; and the membrane cup lure TRE1256 all in combination with the 3.2-mm aperture acetic acid vial were compared in three orchards during Experiment 4: Azwell from 11 – 25 June, Wapato from 30 June to 13 July, and twice in the Moxee orchard from 24 August to 7 September and 31 August to 14 September. Experiment 5 was conducted with the TRE1256 membrane cup lure loaded with 2-phenylethanol and either the TRE1468 membrane cup or the 8.0-ml vial with a 3.2-mm aperture loaded with acetic acid. This test was conducted on two dates in both the Moxee (13 to 27 July and 17 to 31 August) and Ashland (25 July to 8 August and 21 August to 5 September) orchards. Experiment 6 compared traps baited with the membrane cup lure, TRE1468 loaded with acetic acid plus either a 10.0 mg septa (TRE1412) or the TRE1256 membrane cup lure loaded with 2-phenylethanol. This study included 10 replicates in the Moxee and Ashland orchards and was conducted twice from 31 August to 7 September and 21 August to 5 September, respectively. Experiment 7 was conducted twice at the Moxee orchard from 25 July to 8 August and 17 to 31 August, respectively; and in the Ashland orchard from 31 August to 7 September. This experiment included five replicates on each date and compared moth catches in traps baited with the combination of 2-phenylethanol plus acetic acid using either the septa TRE1412 and the membrane cup TRE1468 lures or the membrane cup TRE1256 with the 8.0-ml vial with the 3.2-mm aperture lures.

Two studies compared moth catches in traps baited with either 2-phenylethanol or phenylacetonitrile in combination with acetic acid. Experiment 8 compared moth catches in delta traps baited with the membrane cup lures TRE1256 with 2-phenylethanol or TRE1381 with phenylacetonitrile plus the addition of an 8.0-ml vial with a 3.2-mm aperture loaded with acetic acid. This study was repeated four times during 2016, including from 21 June to 5 July in the Burbank orchard, 16 to 30 August in the Ashland orchard, and 11 to 25 July and 29 August to 12 September in the Moxee orchard. Experiment 9 in 2016 compared moth catches in Multipher traps baited with the 10% sex pheromone lure TRE1229 for *C. rosaceana* alone and in combination with the membrane cup lures TRE1256 or TRE1381. This study was conducted with three replicates of each lure combination in the Tieton, Wapato, and Moxee orchards (N = 9) from 24 July to 21 August.

Moth catches of both *C. pomonella* and *C. rosaceana* in the same trap were recorded in two studies comparing lures used primarily for one species or the other individually or together. Delta traps in Experiment 10 were baited with either the Pherocon CM-DA Combo septa plus the TRE3321 membrane cup lure with acetic acid for *C. pomonella,* the 2-phenylethanol 4.0-ml vial with 1.5-mm aperture lure plus the 8.0-ml vial with a 3.2-mm aperture with acetic acid for *C. rosaceana,* or both sets of lures together. This study was run in both the Wapato and Parker orchards from 9 to 23 July. Experiment 11 was conducted in the Ashland orchard in 2016 comparing moth catches in delta traps baited with the membrane cup lure TRE1468 loaded with acetic acid and either the TRE1225 septa lure loaded with pear ester, DMNT, and codlemone for *C. pomonella,* the membrane cup lure TRE1256 for *C. rosaceana,* or both lures. This study was run from 21 August to 2 September 2016 with 10 replicates.

### Monitoring under mating disruption

Studies were established to examine the effectiveness of 2-phenylethanol plus acetic acid-baited traps for monitoring *C. rosaceana* in orchards treated with and without MD. The size of replicated plots treated with different sex pheromone dispensers in each orchard ranged from 2 - 7 ha. Growers at each study site applied MD dispensers for both *C. pomonella* and *C. rosaceana* during early to mid-April. Four types of MD dispensers were used in these studies and their deployment was randomized in each orchard. During 2015 three replicates of two dispenser treatments (2.0 ha) were established in the Naches orchard: plots were treated with either Cidetrak CMDA Combo PP or a proprietary dispenser Cidetrak CMDA-LR loaded with codlemone, pear ester, and an unspecified four-component blend for leafrollers. Three replicates of two types of dispensers were applied for *C. rosaceana* at the Burbank study site. These two dispensers were either Cidetrak CMDA-LR or the Cidetrak CMDA-LR Meso dispenser. The Cidetrak CMDA-LR Meso dispenser was loaded with 10-times more active material than the former dispenser and was applied at a corresponding 10% rate at 80 ha^−1^. In addition, six plots treated with Isomate CM Flex at 800 ha^−1^ for only *C. pomonella* were also included in this study. During 2016 a similar study was conducted at three sites, and included seven paired replicates (Wapato, N = 3; Tieton, N = 2; and E. Wenatchee, N = 2) of adjacent 4.0-ha plots treated with either Cidetrak CM-DA Meso alone or in combination with a proprietary Cidetrak LR Meso dispenser loaded with (*Z*)-11-tetradecen-1-yl acetate. The two Meso dispensers were paired on plastic clips and applied at 80 ha^−1^.

Study sites were monitored with three different types of lures in delta traps. Traps were baited with either the sex pheromone for *C. rosaceana* (OBLR/W, 3223), the combination lure loaded with pear ester and codlemone (TRE3461) plus the Pherocon AA lure (TRE3321) loaded with acetic acid for *C. pomonella,* or one of several formulations of 2-phenylethanol plus a 8.0-ml vial lure with a 3.2-mm aperture loaded with acetic acid. During 2015, traps with the 2-phenylethanol 4.0-ml vial lure with a 1.5-mm aperture were used in the Naches orchard; and traps with the TRE1256 membrane cup lure loaded with 2-phenylethanol were used in the Burbank orchard. Traps baited with TRE1256 and acetic acid vials were used in all replicates in 2016. Traps for *C. pomonella* were placed at a density of one per ha. Traps with either sex pheromone or 2-phenylethanol and acetic acid lures for *C. rosaceana* were used at a density of one per 2.0-ha. Traps were placed in each orchard in early May and were checked at least once every two weeks in 2015 and once every four weeks in 2016 until mid-September. Sex pheromone septa lures for *C. rosaceana* were replaced every four weeks, and the *C. pomonella* septa lures, the 2-phenylethanol vial and membrane cup lures, and the acetic acid vials were replaced every eight weeks. Adult female *C. rosaceana* caught in traps placed in both the untreated and sex pheromone-treated plots were dissected to determine their mating status. Fruit were sampled during September in both years to detect leafroller larval feeding damage. During 2015 1,200 fruits were examined by sampling thirty fruit from mid-canopy from 40 trees in each plot. A larger fruit sample was taken in 2016 (3,000 fruits per plot), except in the Tieton orchard which was in its third-leaf and not allowed by the farm manager to produce fruit.

### Statistical analyses

Count data were subjected to a square root transformation before analysis of variance (ANOVA) to normalize their variances, and this was measured with a Shapiro-Wilks test. A randomized complete block design was used in most studies with date or orchard as the blocking variable (Statistix 9, Analytical Software, Tallahassee, FL). Treatment means were separated using Tukey’s HSD test with an experiment-wise *α*-value of 0.05 following a significant ANOVA.

## Results

### Lure development

Weight loss over a 14-d period at 25 ° C varied >7-fold among the acetic acid and 2-phenylethanol lures tested in our studies (Table 1). The rate of weight loss for lures used in the seasonal field trials also varied widely over a 12-wk period. For example, the membrane cup lure TRE1256 had a consistent weight loss over this longer time period, slope = 0.0023, *P* = 0.54. The acetic acid vials with 1.0-mm and 3.2-mm apertures clearly had different mean weight losses (Table 1), and the vial with the larger aperture which was the one used in seasonal field trials had a consistent weight loss over a 12-week period at 25 °C, slope = −0.8794, *P* = 0.21. In contrast, the acetic acid membrane cup lure TRE1468 had a significant linear decline in weight loss as a function of time, slope = −4.069, R^2^ = 0.90, *P* < 0.001. After six weeks the mean daily weight loss of the TRE1468 membrane cup lure was reduced to a level (5.09 mg d^−1^) similar to that of a new (< 2-weeks-old) TRE3321 lure (Table 1). Red rubber septa loaded with 10.0 mg of 2-phenylethanol had a relatively short active release phase with only a minimal weight loss (< 0.1 mg d^−1^) measured during the third week of aging at 25 °C. While, the proprietary membrane cup lure loaded with phenylacetonitrile TRE1381 was purported to have the same plastic membrane and to be loaded with the same volume of active compound (800 mg) as the 2-phenylethanol lure (TRE1256); its mean weight loss was nearly 4-fold higher over the initial two-week period of aging (Table 1). TRE1381 also had a consistent weight loss over the extended 12-week assessment period, slope = −0.0218, *P* = 0.27.

**Table 1.**
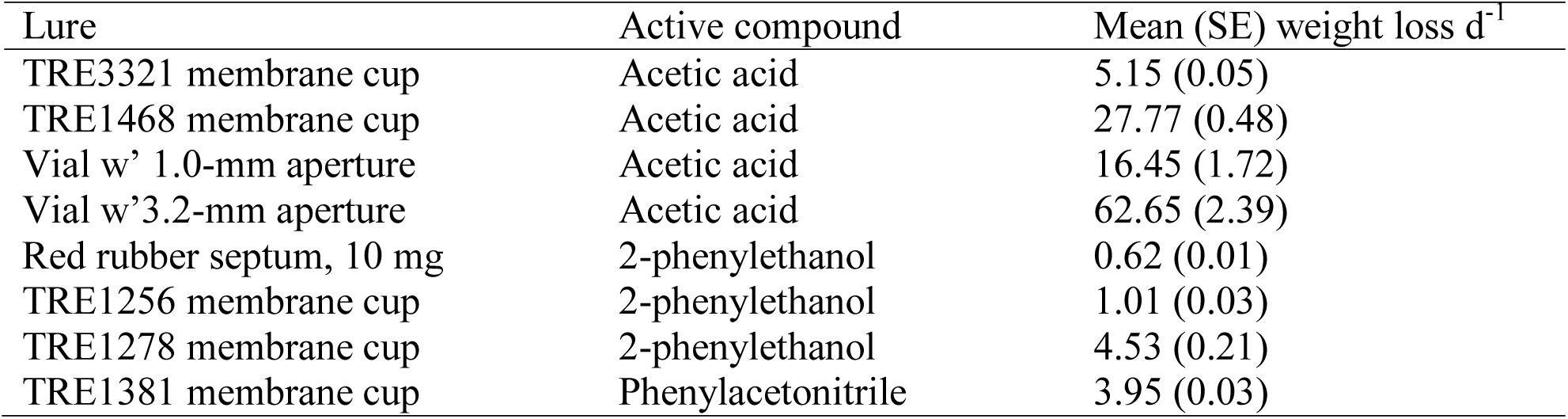
Mean (SE) daily weight loss (mg) at 25 °C over a 14-d period of various lures loaded with acetic acid, 2-phenylethanol, or phenylacetonitrile, N = 5

| Lure | Active compound | Mean (SE) weight loss d<sup>-1</sup> |
| --- | --- | --- |
| TRE3321 membrane cup | Acetic acid | 5.15 (0.05) |
| TRE1468 membrane cup | Acetic acid | 27.77 (0.48) |
| Vial w' 1.0-mm aperture | Acetic acid | 16.45 (1.72) |
| Vial w' 3.2-mm aperture | Acetic acid | 62.65 (2.39) |
| Red rubber septum, 10 mg | 2-phenylethanol | 0.62 (0.01) |
| TRE1256 membrane cup | 2-phenylethanol | 1.01 (0.03) |
| TRE1278 membrane cup | 2-phenylethanol | 4.53 (0.21) |
| TRE1381 membrane cup | Phenylacetonitrile | 3.95 (0.03) |

The type of 2-phenylethanol or acetic acid release device used in delta traps had a significant effect on trap performance across a series of experiments (Table 2). Traps with vials losing higher amounts of acetic acid caught significantly more total moths than traps with the membrane cup lure TRE3321 (Experiment 1). Catch of female moths was significantly higher with the use of the 3.2-mm aperture vial than TRE3321 and the vial with the smaller aperture caught an intermediate number of female moths. Total or female moth catch in Experiment 2 did not vary with the use of either the 1.5-mm aperture vial or the membrane cup TRE1256 lure loaded with 2-phenylethanol when used with the 3.2-mm aperture vial with acetic acid. However, moth catch was significantly reduced in Experiment 3 when traps were baited with the larger membrane cup lure TRE1278 loaded with 2-phenylethanol compared with TRE1256 when used in combination with the 3.2-mm aperture acetic acid vial. Septa lures loaded with either 5.0 or 10.0 mg 2-phenylethanol caught similar number of total moths as traps with TRE1256 when the 3.2-mm aperture vial with acetic acid was used in Experiment 4. All three lures outperformed the 1.0-mg septa lure in this experiment. Differences in female moth catches among lure treatments were more variable with the TRE1256 and 10.0-mg septa lure catching similar numbers and the 5.0-mg septa caught fewer females than the TRE1256-baited traps. No significant difference in total or female moth catch occurred in Experiment 5 between traps baited with TRE1256 and either the membrane cup lure TRE1468 or the vial with a 3.2-mm aperture loaded with acetic acid. Similarly, no significant differences in total or female moth catch occurred in traps baited with TRE1468 loaded with acetic acid and either the 10-mg septa or TRE1256 lures loaded with 2-phenylethanol in Experiment 6. However, the lure combination of TRE1256 plus a vial with a 3.2-mm aperture caught significantly more total moths than traps with a 10-mg septa and TRE1468.The difference in female moth catch between these two lure combinations was not significant, *P* = 0.08.

**Table 2.**
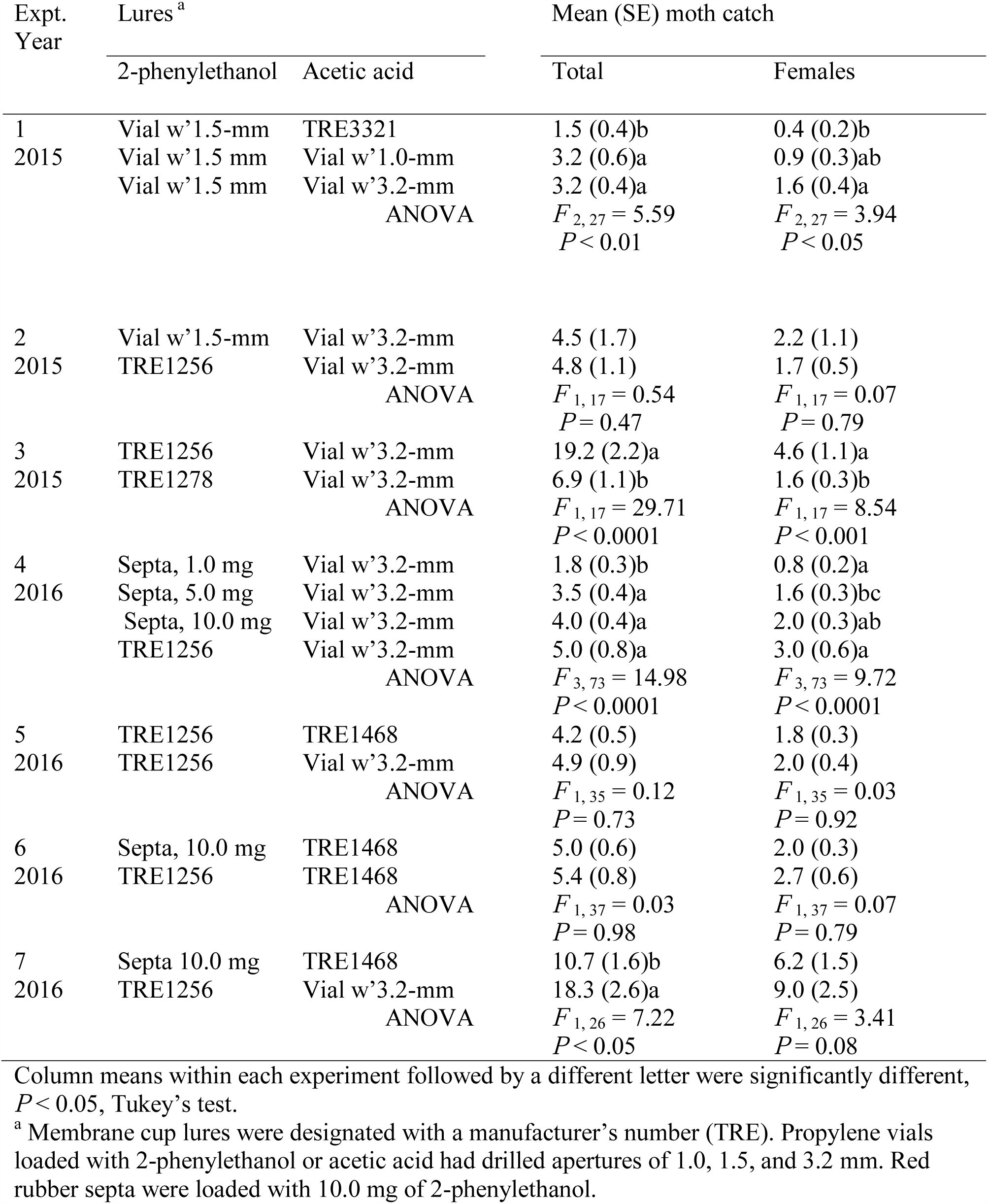
Summary of experiments comparing moth catch of *C. rosaceana* with different combinations of acetic acid (AA) and 2-phenylethanol (PET) lures, 2015-2016

| Expt. | Lures <sup>a</sup> |  | Mean (SE) moth catch |  |
| --- | --- | --- | --- | --- |
| Year | 2-phenylethanol | Acetic acid | Total | Females |
| 1 | Vial w' 1.5-mm | TRE3321 | 1.5 (0.4)b | 0.4 (0.2)b |
| 2015 | Vial w' 1.5 mm | Vial w' 1.0-mm | 3.2 (0.6)a | 0.9 (0.3)ab |
|  | Vial w' 1.5 mm | Vial w' 3.2-mm | 3.2 (0.4)a | 1.6 (0.4)a |
| | ANOVA | | $F_{2,27} = 5.59$ | $F_{2,27} = 3.94$ |
| | | | $P < 0.01$ | $P < 0.05$ |
| 2 | Vial w' 1.5-mm | Vial w' 3.2-mm | 4.5 (1.7) | 2.2 (1.1) |
| 2015 | TRE1256 | Vial w' 3.2-mm | 4.8 (1.1) | 1.7 (0.5) |
| | ANOVA | | $F_{1,17} = 0.54$ | $F_{1,17} = 0.07$ |
| | | | $P = 0.47$ | $P = 0.79$ |
| 3 | TRE1256 | Vial w' 3.2-mm | 19.2 (2.2)a | 4.6 (1.1)a |
| 2015 | TRE1278 | Vial w' 3.2-mm | 6.9 (1.1)b | 1.6 (0.3)b |
| | ANOVA | | $F_{1,17} = 29.71$ | $F_{1,17} = 8.54$ |
| | | | $P < 0.0001$ | $P < 0.001$ |
| 4 | Septa, 1.0 mg | Vial w' 3.2-mm | 1.8 (0.3)b | 0.8 (0.2)a |
| 2016 | Septa, 5.0 mg | Vial w' 3.2-mm | 3.5 (0.4)a | 1.6 (0.3)bc |
|  | Septa, 10.0 mg | Vial w' 3.2-mm | 4.0 (0.4)a | 2.0 (0.3)ab |
|  | TRE1256 | Vial w' 3.2-mm | 5.0 (0.8)a | 3.0 (0.6)a |
| | ANOVA | | $F_{3,73} = 14.98$ | $F_{3,73} = 9.72$ |
| | | | $P < 0.0001$ | $P < 0.0001$ |
| 5 | TRE1256 | TRE1468 | 4.2 (0.5) | 1.8 (0.3) |
| 2016 | TRE1256 | Vial w'3.2-mm | 4.9 (0.9) | 2.0 (0.4) |
| | | ANOVA | $F_{1,35} = 0.12$ | $F_{1,35} = 0.03$ |
| | | | $P = 0.73$ | $P = 0.92$ |
| 6 | Septa, 10.0 mg | TRE1468 | 5.0 (0.6) | 2.0 (0.3) |
| 2016 | TRE1256 | TRE1468 | 5.4 (0.8) | 2.7 (0.6) |
| | | ANOVA | $F_{1,37} = 0.03$ | $F_{1,37} = 0.07$ |
| | | | $P = 0.98$ | $P = 0.79$ |
| 7 | Septa 10.0 mg | TRE1468 | 10.7 (1.6)b | 6.2 (1.5) |
| 2016 | TRE1256 | Vial w'3.2-mm | 18.3 (2.6)a | 9.0 (2.5) |
| | | ANOVA | $F_{1,26} = 7.22$ | $F_{1,26} = 3.41$ |
| | | | $P < 0.05$ | $P = 0.08$ |

Column means within each experiment followed by a different letter were significantly different, $P < 0.05$ , Tukey's test.
<sup>a</sup> Membrane cup lures were designated with a manufacturer's number (TRE). Propylene vials loaded with 2-phenylethanol or acetic acid had drilled apertures of 1.0, 1.5, and 3.2 mm. Red rubber septa were loaded with 10.0 mg of 2-phenylethanol.

Experiments 8 and 9 compared moth catches with membrane cup lures loaded with either 2-phenylethanol or phenylacetonitrile. First, significant differences in both total (*F*_1, 35_ = 4.70, *P* < 0.05) and female (*F* _1, 35_ = 4.95, *P* < 0.05) moth catches were found between traps baited with acetic acid plus either 2-phenylethanol or phenylacetonitrile (Fig. 1). Second, the addition of either 2-phenylethanol or phenylacetonitrile significantly reduced the catch of male moths in Multipher traps baited with sex pheromone, *F* _2, 24_ = 4.62, *P* < 0.05 (Fig. 2).

**Fig. 1.**
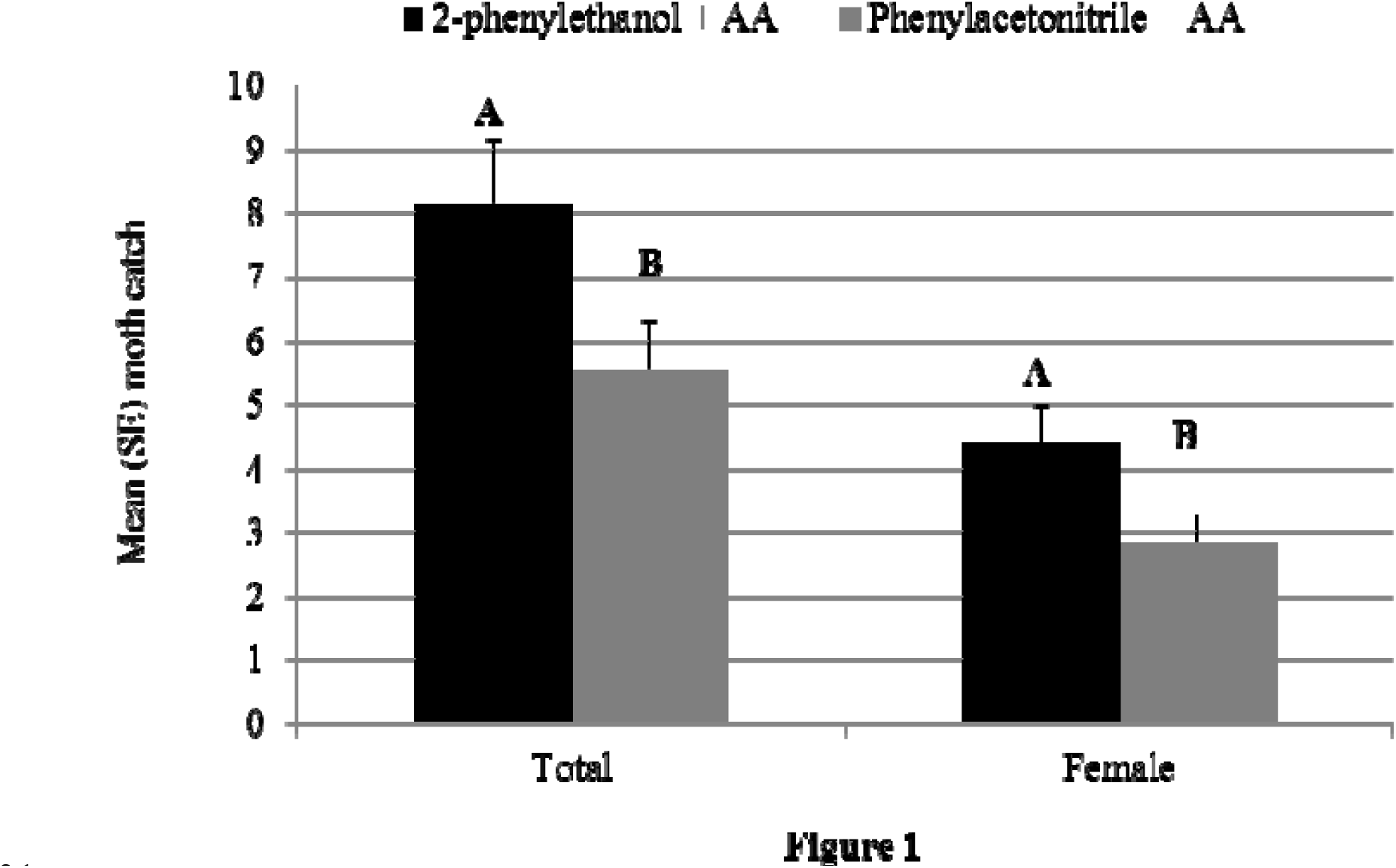
Experiment 8: comparison of mean (SE) total and female moth catches of *C. rosaceana* in delta traps baited with plastic vials with a 3.2-mm aperture in lid loaded with 5.0 ml of acetic acid and proprietary plastic membrane cup lures loaded with either 2-phenylethanol (TRE1256) or phenylacetonitrile (TRE1381).

**Fig. 2.**
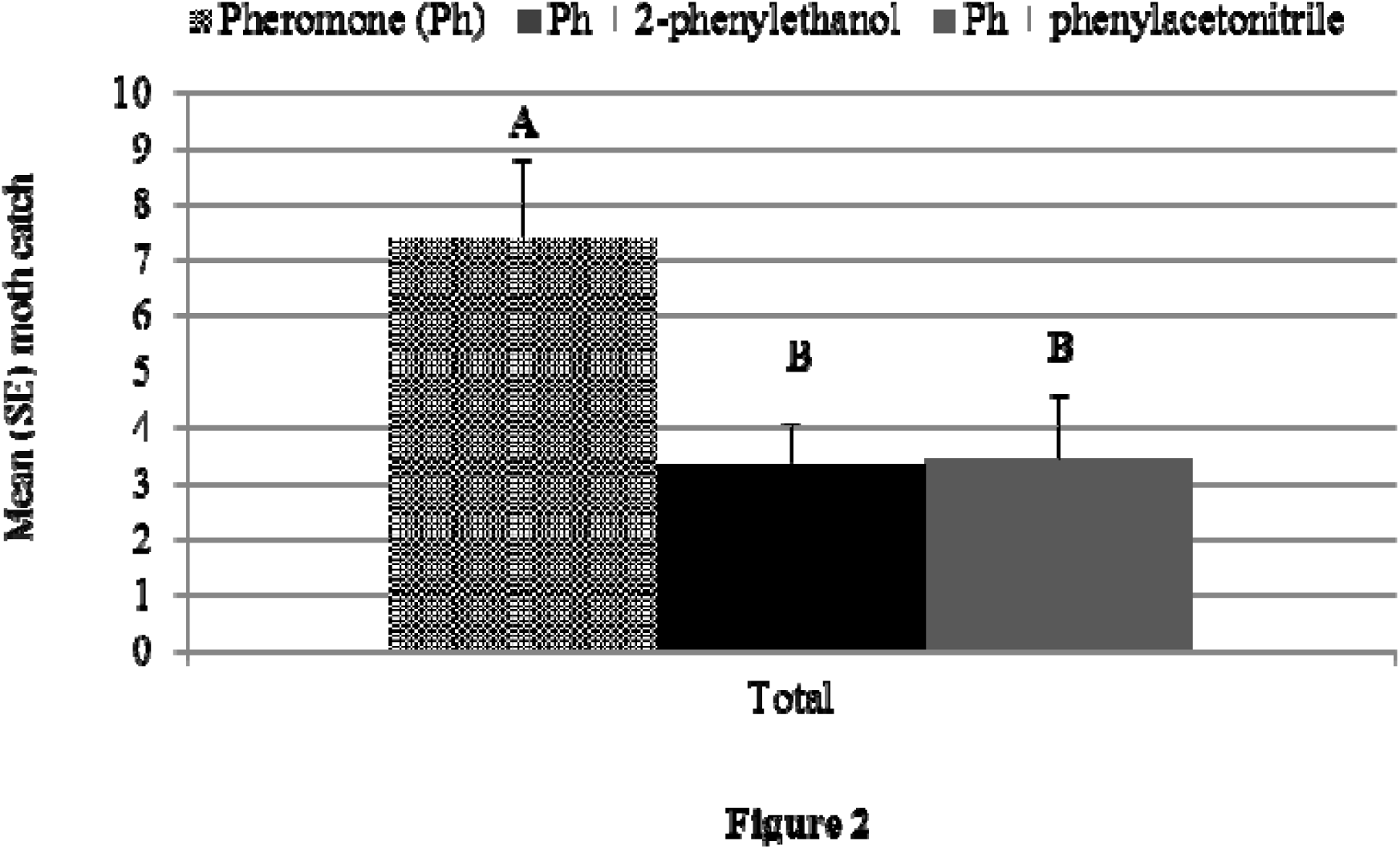
Experiment 9: comparison of mean (SE) male catches of *C. rosaceana* in Multipher 1 plastic bucket traps baited with a red rubber septum loaded with the 4-component sex pheromone (TRE1229), or the same septum plus a proprietary plastic membrane cup lure loaded with 2-phenylethanol (TRE1256) or phenylacetonitrile (TRE1381). Propylene glycol (250 ml) was used in the bottom of these traps to retain moths.

Results were variable in Experiments 10 and 11 for the dual captures of *C. pomonella* and *C. rosaceana* when different lures for each species were combined in traps (Table 3). For example, the addition of the 2-phenylethanol 1.5-mm aperture vial to traps baited with a codlemone and pear ester loaded septum plus the acetic acid 3.2-mm aperture vial did not affect moth catches of *C. pomonella.* Similarly, catch of *C. rosaceana* was not impacted with the two vial lures when the binary septa were added. In contrast, use of a ternary septum lure that included codlemone, pear ester, and DMNT significantly reduced *C. rosaceana* total and female numbers compared with the 2-phenylethanol when used with the membrane cup acetic acid lure TRE1468 (Table 3). This adverse effect from adding DMNT was also seen with the significantly lower catch of *C. rosaceana* that occurred in traps baited with only the ternary septa plus an acetic acid lure.

**Table 3.** Catches of *C. pomonella* and C. *rosaceana* in traps baited with different combinations of lures, including acetic acid (AA), 2-phenylethanol (PET), pear ester (PE), (*E*,*E*)-4,8-dimethyl-1,3,7-nonatriene (DMNT), and codlemone, the sex pheromone (Ph) of *C. pomonella*, 2015–2016

|  |  | Mean (SE) moth catch |  |  |  |
| --- | --- | --- | --- | --- | --- |
| Expt. |  | *C. pomonella* |  | *C. rosaceana* |  |
| Year | Lures <sup>a</sup> | Total | Female | Total | Female |
| 10 | Ph-PE septum + AA vial | 34.1 (5.0)a | 13.4 (2.2)a | 10.4 (2.4)b | 2.6 (0.6)b |
| 2015 | PET vial + AA vial | 2.5 (0.5)b | 2.0 (0.4)b | 35.8 (9.3)a | 8.7 (2.3)a |
|  | Ph-PE septum + PET and AA vials | 30.8 (4.1)a | 10.0 (1.8)a | 29.4 (6.6)a | 7.5 (1.2)a |
| | ANOVA | $F_{2,26} = 55.27$ | $F_{2,26} = 11.22$ | $F_{2,26} = 8.03$ | $F_{2,26} = 7.77$ |
| | | $P < 0.0001$ | $P < 0.001$ | $P < 0.01$ | $P < 0.01$ |
| 11 | Ph-DMNT-PE septum + AA membrane | 20.2 (4.1)a | 7.1 (1.6)a | 0.3 (0.2)c | 0.1 (0.1)c |
| 2016 | PET and AA membranes | 0.6 (0.2)b | 0.2 (0.1)b | 9.2 (1.5)a | 4.6 (0.8)a |
|  | Ph-DMNT-PE septum + PET and AA membrane | 18.4 (3.5)a | 5.7 (0.8)a | 2.1 (0.6)b | 1.2 (0.4)b |
| | ANOVA | $F_{2,27} = 34.45$ | $F_{2,27} = 29.48$ | $F_{2,27} = 37.67$ | $F_{2,27} = 33.69$ |
| | | $P < 0.0001$ | $P < 0.0001$ | $P < 0.0001$ | $P < 0.0001$ |

Column means within each experiment followed by a different letter were significantly different, $P < 0.05$ , Tukey's test.
<sup>a</sup> Lures included grey halobutyl septa loaded with either pear ester and codlemone (Ph-PE, TRE3461) or a combination of DMNT, pear ester, and codlemone (Ph-DMNT-PE, TRE1225); 8-ml vials with a 3.2-mm aperture loaded with 5.0 ml of acetic acid, and 4.0-ml vials with a 1.5-mm aperture loaded with 3.0 ml of 2-phenylethanol; and membrane cup lures loaded with either acetic acid (TRE1468) or 2-phenylethanol (TRE1256).

### Monitoring under mating disruption

Similar and significant differences in catches of *C. rosaceana* were found among lures in orchard plots treated with MD dispensers in both study years (Table 4). A significant interaction of MD dispenser treatment × lure type was found in the ANOVA for total moth catches in both years. Moth catch was highest in sex pheromone-baited traps placed in untreated plots. Total moth catch in traps baited with 2-phenylethanol and acetic acid in both untreated and treated plots had the second highest total moth catch, and moth catch in either the sex pheromone-baited traps placed in MD-treated plots or catch in traps baited with codlemone, pear ester and the membrane cup lure TRE3321 in either the untreated or MD-treated plots were the lowest. Cumulative mean female moth catch were similar and significantly higher in both the untreated and MD-treated plots then in traps baited with the codlemone plus pear ester septa and the membrane cup TRE3321 acetic acid lure. Few unmated *C. rosaceana* females were caught in traps baited with 2-phenylethanol and acetic acid in untreated and MD-treated plots in either year: 18 and 22% in 2015 and 5 and 10% in 2016, respectively. No sampled fruit from any orchard in either year were found to have leafroller larval injury.

**Table 4.** Catches of oblique banded leafroller (OBLR), *C. rosaceana* in traps baited with either sex pheromone (OBLR PH), 2-phenylethanol (PET) plus acetic acid (AA) or the sex pheromone of *C. pomonella* and pear ester (CM Ph-PE) plus AA in season-long trials conducted in six and seven paired apple blocks treated with or without sex pheromone mating disruption (MD) for *C. rosaceana* in Washington State during 2015 and 2016, respectively

| Year | Dispenser<br>treatment <sup>a</sup> | Mean (SE) total moth catch per trap |  |  | Mean (SE) female moth catch per trap |  |
| --- | --- | --- | --- | --- | --- | --- |
|  |  | OBLR PH | PET + AA<sup>b</sup> | CM Ph-PE + AA<sup>c</sup> | PET + AA<sup>b</sup> | CM Ph-PE + AA<sup>c</sup> |
| 2015 | None | 51.9 (7.0)a | 13.3 (2.2)b | 2.0 (0.7)c | 4.3 (0.9)A | 0.6 (0.2)B |
|  | OBLR MD | 0.1 (0.1)c | 11.2 (1.7)b | 2.1 (1.0)c | 3.0 (0.8)A | 0.7 (0.3)B |
| | | Treatment: | $F_{1,42} = 91.40, P < 0.0001$ | | Treatment: | $F_{1,32} = 0.93, P = 0.35$ |
| | ANOVA | Lure: | $F_{2,42} = 38.79, P < 0.0001$ | | Lure: | $F_{1,32} = 32.15, P < 0.0001$ |
| | | Treatment * Lure: | $F_{2,42} = 75.21, P < 0.0001$ | | Treatment * Lure: | $F_{1,32} = 1.38, P = 0.25$ |
| 2016 | None | 59.1 (19.7)a | 18.3 (7.2)b | 1.0 (0.7)c | 5.3 (1.7)A | 0.6 (0.6)B |
|  | OBLR MD | 1.0 (0.4)c | 14.4 (4.1)b | 0.4 (0.3)c | 5.7 (1.5)A | 0.1 (0.1)B |
| | | Treatment: | $F_{1,36} = 13.24, P < 0.001$ | | Treatment: | $F_{1,24} = 0.00, P = 0.99$ |
| | ANOVA | Lure: | $F_{2,36} = 12.72, P < 0.001$ | | Lure: | $F_{1,24} = 28.11, P < 0.0001$ |
| | | Treatment * Lure: | $F_{2,36} = 11.26, P < 0.001$ | | Treatment * Lure: | $F_{1,24} = 0.16, P = 0.70$ |

Mean total catch followed by a different lowercase letter and mean female catch followed by a different uppercase letter within each year were significantly different, $P < 0.05$ , Tukey's test.
<sup>a</sup> Dispensers used in these studies included: Cidetrak CMDA Combo and Isomate Flex (None) or Cidetrak CMDA-LR Combo or Cidetrak CMDA-LR Combo Meso dispensers (OBLR MD) in 2015, and Cidetrak CMDA Combo Meso (None) and Cidetrak CMDA Combo Meso plus Combo LR Meso dispensers in 2016. All Meso dispensers were applied at 80 dispensers ha<sup>-1</sup> and other dispensers were applied at 800 ha<sup>-1</sup>
<sup>b</sup> Traps were baited with two different 2-phenylethanol lures in combination with a acetic acid lure (8.0-ml vial with 3.2-mm aperture). These included 4.0-ml vials with 1.5-mm apertures loaded with 2-phenylethanol in 2015 and the membrane cup lure TRE1256 in 2016.
<sup>c</sup> Traps were baited with TRE3461 septa lures loaded with codlemone and pear ester and membrane cup TRE3321 lures loaded with acetic acid.

## Discussion

The combined use of 2-phenylethanol and acetic acid was shown to be an effective lure to monitor *C. rosaceana* within orchards treated with sex pheromones for MD. In comparison, sex pheromone-baited traps within MD-treated orchards caught almost no moths during the season as has been previously shown with a range of MD technologies, including, sprayables, hand-applied dispensers, and aerosols (Knight et al. 1998; Evenden et al. 1999; Knight 2000; Judd and Gardiner 2008). Moth catch was also 5 to 40-times higher with the new binary lure than in traps baited with pear ester and acetic acid. Previously, the pear ester plus acetic acid lure combination was found to provide some useful information in regards to pest density of both *C. rosaceana* and *P. pyrusana* (Knight et al. 2014). Unfortunately, no leafroller fruit injury was detected in the various orchards monitored in 2015–2016, and correlations of moth catches with this new binary lure and larval pest pressure was not possible. Future studies will survey orchards with a range of pest pressures of *C. rosaceana* to establish whether useful action thresholds can be established with the use of a binary 2-phenylethanol and acetic acid lure.

Our array of field experiments have helped to optimize the emission characteristics needed for both 2-phenylethanol and acetic acid lures, i.e. ease in handling, attractiveness, and longevity. The significant reduction in moth catch that occurred with the membrane cup lure TRE1278 compared with TRE1256 defined the upper limit of the emission rate of 2-phenylethanol. A rubber septum loaded with 10.0 mg of 2-phenylethanol was effective in catching moths, but was too short lived. In comparison, the membrane cup TRE1256 lure provided a high level of release over a long time period. The membrane cup lure TRE1468 provided a level of emission of acetic acid similar to the 1.0-mm aperture vial for a number of weeks, but further refinement of this lure (characteristics of the plastic membrane and initial loading) is still needed to achieve a somewhat higher emission rate over a longer time period.

We found that the combination of 2-phenylethanol and acetic acid was more attractive for *C. rosaceana* than the blend of phenylacetonitrile and acetic acid. Conversely, the reverse in the attractiveness of these two binary blends was found in Canada for the eye-spotted bud moth, *Spilonota ocellana* (Denis & Schiffermüller) (El-Sayed et al. 2016). Studies in Sweden found that both sexes of a number of tortricids, including *P. heparana, P. cerasana* (Hübner), *Archips rosana* (L.), *A. podana* Scopoli, *S. ocellana,* and *Hedya nubiferana* Haworth, occurred in traps baited with acetic acid and either 2-phenylethanol or phenylacetonitrile, but the attractiveness of the two lures was only compared with the two *Pandemis* species in this study, and this did not differ (Giacomuzzi et al. 2016).

Unfortunately, the combination of phenylacetonitrile and acetic acid was found to reduce catches of *C. pomonella* when used in traps with pear ester lures (Knight et al. 2016). The effectiveness of codlemone, pear ester, and acetic acid used together was not affected by the inclusion of either phenylacetonitrile (Knight et al. 2016) or 2-phenylethanol (Fig. 1). Catches of *C. rosaceana* were reduced in traps when DMNT was added to a septum loaded with codlemone and pear ester, and this is significant because the addition of DMNT has been shown to improve monitoring for *C. pomonella*; especially within orchards treated with dispensers containing both codlemone and pear ester (Knight et al. 2015). Thus, additional studies should assess the optimal lure needed to monitor the diverse tortricid pest complexes which occur among geographical regions. This will include the use of blends of varying ratios of 2-phenylethanol and phenylacetonitrile, and perhaps the addition of other lure components, such as benzyl alcohol (Giacomuzzi et al. 2016). The use of compatible multi-component lures placed in single traps may reduce management costs and be conducive to the development of smart-trap technologies for tortricid pests (Kim et al. 2011).

Moth catches of *C. rosaceana* in non-disrupted orchards were significantly lower than with sex pheromone-baited traps. This result was different from that found with phenylacetonitrile plus acetic acid compared with a sex pheromone lure for *P. pyrusana* (Knight et al. 2016). One concern may be that the 2-phenylethanol plus acetic acid lure might not be attractive enough to allow managers to track the seasonal phenology of *C. rosaceana* in MD-treated orchards. With *C. pomonella* the addition of pear ester to codlemone significantly increased the effectiveness of monitoring in MD-treated orchards (Knight et al. 2005). Similar additive results have been reported with host plant volatiles and the sex pheromones of other important tortricid species, such as *Eupoecilia ambiguella* Hübner, *Grapholita molesta* (Büsck), and *Lobesia botrana* (Schmidt-Büsser et al. 2009; Varela et al. 2011; von Arx et al. 2012). Interestingly, the use of acetic acid with either the sex pheromone or these host plant volatiles has not been reported. Yet, the interaction of host volatiles and species’ sex pheromone can have inhibitory or repellent effects (Reddy and Guerrero 2004). Here, we found that the addition of either aromatic volatile to the sex pheromone of *C. rosaceana* significantly reduced male moth catch. Further tests should evaluate whether the addition of acetic acid with 2-phenylethanol can override this behavioral effect, and also measure any effects of combining sex pheromone with the aromatic and acetic acid lures on the catch of female moths in traps. These data will be important prerequisites in order to establish action thresholds for supplementary management actions based on bisexual moth catches of *C. rosaceana* in MD-disrupted orchards.

## Acknowledgements

We would like to thank Duane Larson and Chey Temple, Agricultural Research Service, Wapato, WA for their technical assistance in the laboratory and field. We would like to acknowledge our appreciation to Bill Lingren, Trécé Inc., Adair, OK for donating many of the lures used in our studies. We also thank Carolina Oyarzún, Instituto de Producción y Sanidad Vegetal, Facultad de Ciencias Agrarias, Universidad Austral de Chile, Valdivia, Chile; and Valentina Mujica, Instituto Nacional de Investigación Agropecuaria, National Research Program in Fruit Production, Canelones, Uruguay for their help in sexing and dissecting moths. This project was supported with partial funding from the Washington Tree Fruit Research Commission, Wenatchee, WA.

## References

Alway T, 2003. Exploring alternatives to organophosphate insecticides with apple and pear: the Areawide II Project in Washington State. Compact Fruit Tree 36, 91–93.

Barnett WW, Bentley WJ, Bethell RS, Pickel C, Weddle PW, Zalom FG, 1991. Managing pests in apples and pears. In: Integrated pest management of apples and pears. Ed. by Flint ML, University of California Press, Oakland, CA, 21–53.

Becher PG, Bengtsson M, Hansson BS, Witzgall P, 2010. Flying the fly: long-range flight behavior of *Drosophila melanogaster* to attractive odors. J Chem Ecol, 36, 599–607.

Beers EH, Brunner JF, Willett MJ, Warner GW, 1993. Orchard pest management: a resource book for the Pacific Northwest. Good Fruit Grower, Yakima, WA.

Bengtsson M, Jaastad G, Knudsen G, Kobro S, Bäckman AC, Pettersson E, Witzgall P, 2006. Plant volatiles mediate attraction to host and non-host plants in apple fruit moth, *Argyresthia conjugella*. Entomol Exper Appl, 118, 77–85.

Boevé JL, Lengwiler U, Tollsten L, Dorn S, Turlings CJ, 1996. Volatiles emitted by apple fruitlets infested by larvae of the European apple sawfly. Phytochem, 42, 373–381.

Bruce TJA, Wadhams LJ, Woodcock CM, 2005. Insect host location: a volatile situation. Trends Plant Sci, 10, 269–274.

Brunner JF, Beers EH, 1990. Pandemis and obliquebanded leafrollers. Washington State Univ Extension Bull, 1582, Wenatchee, WA. 6 pp.

Buchbauer G, Jirovetz L, Wasicky M, Nikiforov A, 1993. Headspace and essential oil analysis of apple flowers. J Agric Food Chem, 41, 116–118.

Chapman PJ, 1973. Bionomics of the apple-feeding Tortricidae. Annu Rev Entomol, 18, 73–96.

Cossentine JE, Jensen LB, Deglow EK, 2003. Strategy for orchard use of *Bacillus thuringiensis* while minimizing impact on *Choristoneura rosaceana* parasitoids. Entomol Exper Appl, 109, 205–210.

Curkovic T, Brunner JF, 2005. Residual and sub-lethal effects of an attracticide formulation on *Choristoneura rosaceana* (Harris), *Pandemis pyrusana* Kearfott, and *Cydia pomonella* (L.) males (Lepidoptera: Tortricidae). Crop Protec, 24, 637–641.

Dunley JE, Brunner JF, Doerr MD, Beers EH, 2006. Resistance and cross-resistance in populations of leafrollers, *Choristoneura rosaceana* and *Pandemis pyrusana* in Washington apples. J Insect Sci, 6, 1–14.

El-Sayed AM, Knight AL, Byers JA, Judd GJR, Suckling DM, 2016. Caterpillar-induced plant volatiles attract conspecific adults in nature. Sci Reports (in press).

Evenden ML, Judd GJR, Borden JH, 1999. Mating disruption of two sympatric, orchardinhabiting tortricids, *Choristoneura rosaceana* and *Pandemis limitata* (Lepidoptera: Tortricidae) with pheromone components of both species blends. J Econ Entomol, 92, 380–390.

Giacomuzzi V, Abraham J, Angeli S, 2013. Feeding damage of *Pandemis heparana* induces the release of specific volatile compounds from apple plants. Proc Conf Intern Res Food Security, Univ Hohenheim. 17–19 September 2013

Giacomuzzi V, Cappellin L, Khomenko I, Biasioli F, Schutz S, Tasin M, Knight AL, Angeli S, 2016. Emission volatile compounds from apple plants infested with *Pandemis heparana* larvae, antennal response of conspecific adults, and preliminary field trial. J Chem Ecol, (in press).

Jones VP, Eastburn CC, Wilburn TD, Brunner JF, 2005. Instar-specific phenology of *Pandemis pyrusana* and *Choristoneura rosaceana* (Lepidoptera: Tortricidae) in Washington apple orchards. J Econ Entomol, 98, 875–883.

Jones VP, Horton DR, Mills NJ, Unruh TR, Baker CC, Melton TD, Milickzy E, Steffan SA, Shearer PW, Amarasekare KG, 2015. Evaluating plant volatiles for monitoring natural enemies in apple, pear, and walnut orchards. Bio Contr, 102, 53–62.

Judd GJR, Gardiner MGT, 2008. Efficacy of Isomate-CM/LR for management of leafrollers by mating disruption in organic apple orchards of Western Canada. J Entomol Soc British Columbia, 105, 45–60.

Kim Y, Jung S, Kim Y, Lee Y, 2011. Real-time monitoring of oriental fruit moth, *Grapholita molesta,* populations using a remote sensing pheromone trap in apple orchards. J Asia- Pacific Entomol, 14, 259–262.

Knight AL, 2000. Applying sex pheromones – from puffes to air-blast sprayers. Proc WA Hort Assoc, 96, 161–163.

Knight AL, 2001. Monitoring the seasonal population density of *Pandemis pyrusana* (Lepidoptera: Tortricidae) within a diverse fruit crop production area in the Yakima Valley, WA. J Entomol Soc British Columbia, 98, 217–225.

Knight A, 2010. Organic codling moth management in Washington State and the world. Proc WA State Hort Assoc, 106, 131–136.

Knight AL, Thomson DR, Cockfield SD, 1998. Developing mating disruption of obliquebanded leafroller (Lepidoptera: Tortricidae) in Washington State. Environ Entomol, 27, 1080–1088.

Knight AL, Hilton R, Basoalto E, Stelinski LL, 2014. Use of glacial acetic acid to enhance bisexual monitoring of tortricid pests with kairomone lures in pome fruits. Environ Entomol, 43, 1628–1640.

Knight AL, Basoalto E, Katalin J, El-Sayed AM, 2015. A binary host plant volatile lure combined with acetic acid to monitor codling moth (Lepidoptera: Tortricidae). Environ Entomol, 44, 1434–1440.

Knight AL, Basoalto E, Judd GJR, Suckling DM, El-Sayed AM, 2016. Use of phenylacetonitrile plus acetic acid for combined monitoring of *Pandemis pyrusana* and *Cydia pomonella* (Lepidoptera: Tortricidae) in apple orchards. Environ Entomol, (in press).

Knudsen JT, Tollsten L, Bergstron LG, 1993. Floral scents – a checklist of volatile compounds isolated by head-space techniques. Phytochem, 33, 253–280.

Madsen HF, Madsen BJ, 1980. Response of four leafroller species (Lepidoptera: Tortricidae) to sex attractants in British Columbia orchards. Can Entomol, 112, 427–430.

Omata A, Yomogida K, Nakamura S, Hashimoto S, Koba S, Furukawa K, Noro S, 1990. Volatile components of apple flowers. Flavour Fragance J, 5, 19–22.

Pree DJ, Whitty KJ, Pogoda MK, Bittner LA, 2001. Occurrence of resistance to insecticides in populations of the obliquebanded leafroller from orchards. Can Entomol, 133, 93–103.

Reddy GVP, Guerrero G, 2004. Interactions of insect pheromones and plant semiochemicals. Trends Plant Sci, 9, 253–261.

Schmidt-Busser D, von Arx M, Guerin PM, 2009. Host plant volatiles serve to increase the response of male European grape berry moths, *Eupoecilia ambiguella*, to their sex pheromone. J Comp Physiol, 195, 853–864.

Sial AA, Brunner JF, Doerr MD, 2010. Susceptibility of *Choristoneura rosaceana* (Lepidoptera: Tortricidae) to two new reduced-risk insecticides. J Econ Entomol, 103, 140–146.

Smirle MJ, Lowery T, Zurowski CL, 2003. Susceptibility of leafrollers (Lepidoptera: Tortricidae) from organic and conventional orchards to azinphosmethyl, spinosad, and *Bacillus thuringiensis*. J Econ Entomol, 96, 879–884.

Suckling DM, Twidle AM, Gibb AR, Manning LM, Mitchell VJ, Sullivan TES, Wee SL, El- Sayed AM, 2012. Volatiles from apple trees infested with light brown apple moth larvae attract the parasitoid *Dolichogenida tasmanica*. J Agric Food Chem, 60, 9562–9566.

Tasin M, Betta E, Carlin S, Gasperi F, Mattivi F, Pertot I, 2011. Volatiles that encode host-plant quality in the grapevine moth. Phytochem, 72, 1999–2005.

Thayer GR, Gut LJ, Brunner JF, 1997. The active space of leafroller pheromone traps baited with lures of different strength. Proc. Western Orchard Pest & Disease Manage Conf, 71, 14.

Toth M, Szentkiralyi F, Vuts J, Letardi A, Tabilio MR, Jaastad G, Knudsen GK, 2009. Optimization of a phenylacetaldehyde attractant for common green lacewings *(Chrysoperla carnea* s.l.). J Chem Ecol, 35, 449–458.

Varela N, Avilla J, Anton S, Gemeno C, 2011. Synergism of pheromone and host-plant volatile blends in the attraction of *Grapholita molesta* males. Entomol Exper Appl, 141, 114–122.

von Arx M, Schmidt-Busser D, Guerin, 2012, Plant volatiles enhance behavioral responses of grapevine moth males, *Lobesia botrana* to sex pheromone. J Chem Ecol, 38, 222–225.

Zhu J, Park KC, Baker TC, 2003. Identification of odors from overripe mango that attract vinegar flies, *Drosophila melanogaster*. J Chem Ecol, 29, 899–909.

Zhu J, Obrycki JJ, Ochieng SA, Baker TC, Pickett JA, Smiley D, 2005. Attraction of two lacewing species to volatiles produced by host plants and aphid prey. Naturwissen, 92, 277–281.

